# Sperm quality parameters are increased and asymmetric in house mouse hybrids

**DOI:** 10.1101/666511

**Authors:** Iva Martincová, Ľudovít Ďureje, Stuart J. E. Baird, Jaroslav Piálek

## Abstract

Spermatogenesis is a tuned cascade of processes producing sperm; impairment of any phase of this process can affect fitness of males. The level of impairment can be pronounced in hybrids between genetically divergent populations. To explore the effect of hybridization on sperm quality we produced F1 hybrids from 29 wild derived strains of two house mouse subspecies, *M. m. musculus* and *M. m. domesticus*, which diverged 0.5 MY ago. The measured sperm quality traits did not significantly differ between intrasubspecific crosses. Effects of intersubspecific hybridization were dependent on sperm trait and cross direction. The proportion of sperm head abnormalities was increased in F1 intersubspecific hybrids. The frequency of dissociated sperm heads was increased in the *M. m. musculus* × *M. m. domesticus* (♀×♂) F1 but decreased in *M. m. domesticus* × *M. m. musculus* (♀×♂) F1 hybrids, with the difference in medians being more than 180%. We deduce that the dissociated sperm heads trait is associated with the X chromosome and modulated by interaction with the Y chromosome; nevertheless, the high proportion of unexplained variance (55.46 %) suggests the presence of polymorphic autosomal interactions. The reported differences in sperm quality between cross types may be highly relevant to male fitness in zones of secondary contact between the two subspecies. The cross direction asymmetry in frequency of dissociated sperm heads should favour the *M. m. musculus* Y chromosome. This is consistent with the spread of the *M. m. musculus* Y chromosome in nature across the hybrid zone between these two subspecies.

## Introduction

Lower fertility in males due to abnormal spermatozoa has been reported in many animals including humans, but its pathogenic causes, including genetic factors, remain largely unknown (Chen et al. 2016). Data from knockout mice suggest that numerous genes can impair spermiogenesis (reviewed in Chen et al. 2016). In nature, the incidence of genetically induced abnormalities can be affected due to hybridization between diverged populations in zones of secondary contact (Alund et al. 2013). Decreased sperm quality will tend to reduce, and improved sperm quality increase, the spread of causal genes. The amplitude of abnormal sperm morphology changes will be modulated by geographic standing variation, affecting gene flow at local scales. Nevertheless, in some traits hybridization may reveal species-specific effects, affecting gene flow at wider scales along contact zones.

The house mouse hybrid zone (HMHZ) between *Mus musculus musculus* and *M. m. domesticus* whose genomes diverged 0.5 MY ago is among the best studied animal hybrid zones (Baird & Macholán 2012). The zone stretches across Europe from Norway to Bulgaria; more than 2500 km long, itis only 10-20 km wide. The enigmatic behaviour of the sex chromosomes has been especially challenging to our understanding of the extent and causality of introgressive hybridization (Albrechtová et al. 2012). While introgression of the X chromosome is limited in comparison with the autosomal loci Y introgression seems widespread along an 850 km of the contact in Central Europe, and highly asymmetric: Y^musculus^ invades the *domesticus* background, the converse being extremely rare (Macholán et al. 2007, 2008, Ďureje et al. 2012).

To explore the conditions of this Y chromosome spread in a wider context we earlier derived 31 recombinant lines from eight wild-derived strains representing four localities within the two mouse subspecies (Martincová et al. 2019). These strains were first crossed within either subspecies and subsequently reciprocally crossed to raise intersubspecific hybrids. The resulting F1 hybrid males were scored for five phenotypic traits associated with male fitness. Hierarchical analysis of reproductive traits in F1 hybrids with recombined intrasubspecific genomes revealed that body and testis weights and sperm count differed on local scales (i.e. between localities and strains within subspecies) but not at the intersubspecific level (Martincová et al. 2019). On the other hand, two sperm quality traits (frequency of abnormal sperm heads (ASH) and frequency of dissociated sperm heads from tail (DSH)) displayed Y-associated differentiation at the intersubspecific level. In both cases the F1 hybrids with Y^musculus^ performed better than the F1 hybrids possessing Y^domesticus^. However, hierarchical model analysis suggested that the signal detected at the intersubspecific level in ASH might be spurious, rather caused by interactions between one specific Y^domesticus^ haplotype and the X or autosomes in recombinant lines.

In this paper we explore the replicability and generality of the sperm quality observations for 8 Y haplotypes reported by (Martincová et al. 2019). Increasing sample size such that representative genetic variation is captured within each of the subspecies and inter- and intrasubspecific F1 hybrid comparisons we address two questions: (1) Are the proportions of deformed sperm in intersubspecific hybrids different in comparison with intrasubspecific F1 hybrids? (2) Is the sperm quality in intersubspecific F1 hybrids dependent on cross direction and, if so, are frequencies of DSH and ASH decreased in the *domesticus* × *musculus* (♀ × ♂) male progeny when compared with the *musculus* × *domesticus* (♀ × ♂) male progeny? Such a comparative study of the same set of spermrelated traits across different experimental replicates tests the repeatability of subspecies-specific phenotypic variation observations.

## Methods

### Animals

The 29 wild-derived strains used to produce F1 hybrids are maintained in the breeding facility of the Institute of Vertebrate Biology in Studenec (Piálek et al. 2008; licences for keeping animals and experimental work 61974/2017-MZE-17214 and 62065/2017-MZE-17214, respectively). Fourteen of them were derived at University of Montpellier by Annie Orth and François Bonhomme and donated to Studenec in 2017 and 2018. PWD was derived in the Institute of Molecular Genetics (Gregorová & Forejt 2000). Sixteen strains represent sampling of the wild standing variation of *domesticus* genomes and 13 strains *musculus* genomes. Their distribution captures a wealth of genetic variation present within a geographic area covering almost 8.5 M km^2^. Detailed information on the origin of wild-derived strains and numbers of mated sires and dams used are listed in Table 1 and can also be retrieved at https://housemice.cz/. Numbers of sires used to generate intra- and/or intersubspecific hybrids ranged between 1-18 (mean ± SD: 6.14 ± 4.69) and the numbers of dams ranged between 1-15 (7.00 ± 4.49). In total, we scored 178 unique F1 males representing 34 *domesticus* × *domesticus*, 61 *domesticus* × *musculus*,42 *musculus* × *domesticus* and 42 *musculus* × *musculus* crosses. Note, in all notifications of crosses, the female precedes the male (♀ × ♂). Each male was derived from a unique pair of parents and their data are considered independent.

**Table 1.** List of wild-derived strains and numbers of parents used in this study

| Genome | State | WDS | Latitude | Longitude | Sires | Dams |
| --- | --- | --- | --- | --- | --- | --- |
| domesticus | Israel | BIK | 32.766 | 34.960 | 7 | 5 |
| domesticus | Algeria | BZO | 35.698 | -0.634 | 2 | 4 |
| domesticus | Cyprus | DCA | 34.583 | 32.970 | 1 | 0 |
| domesticus | Cyprus | DCP | 34.772 | 32.430 | 8 | 6 |
| domesticus | Denmark | DDO | 55.409 | 9.383 | 2 | 4 |
| domesticus | Georgia | DGA | 41.601 | 42.069 | 4 | 5 |
| domesticus | Israel | DIK | 32.980 | 35.808 | 1 | 0 |
| domesticus | Italy | DJO | 45.527 | 8.542 | 2 | 1 |
| domesticus | Tahiti | DOT | -17.538 | -149.559 | 1 | 0 |
| domesticus | Germany | SCHUNT | 50.427 | 9.598 | 10 | 15 |
| domesticus | UK | SIN | 59.297 | -2.554 | 3 | 4 |
| domesticus | UK | SIT | 59.297 | -2.554 | 8 | 12 |
| domesticus | Germany | STAIL | 50.427 | 9.598 | 2 | 5 |
| domesticus | Germany | STRA | 50.181 | 11.763 | 10 | 14 |
| domesticus | Germany | STRB | 50.181 | 11.763 | 8 | 10 |
| domesticus | France | WLA | 43.605 | 1.444 | 6 | 4 |
| musculus | Czechia | BULS | 50.222 | 13.376 | 12 | 15 |
| musculus | Czechia | BUSNA | 50.228 | 13.370 | 9 | 9 |
| musculus | Armenia | MAM | 38.902 | 46.247 | 1 | 1 |
| musculus | Georgia | MGA | 41.717 | 45.978 | 2 | 3 |
| musculus | Hungary | MHB | 47.495 | 19.040 | 1 | 1 |
| musculus | Poland | MPB | 52.701 | 23.875 | 7 | 8 |
| musculus | Czechia | PWD | 50.013 | 14.481 | 18 | 14 |
| musculus | Slovakia | SENK | 48.299 | 17.349 | 5 | 2 |
| musculus | Estonia | SKA | 58.955 | 24.834 | 7 | 12 |
| musculus | Estonia | SKE | 58.941 | 24.908 | 6 | 6 |
| musculus | Czechia | STUF | 49.200 | 16.064 | 15 | 7 |
| musculus | Czechia | STUS | 49.200 | 16.064 | 14 | 5 |

### Phenotyping

All males were sacrificed by cervical dislocation and dissected at 60 days of age. Spermatozoa were released from the whole left epididymis and transferred into a watch glass with 2ml of 1% sodium citrate (for details see Vyskočilová et al. 2005). Sperm quality parameters were evaluated in a Bürker haemocytometer using an Olympus CX41 microscope under 200× magnification (for details see Vyskočilová et al. (2005)). The frequency of dissociated sperm heads (DSH) was estimated from five haemocytometer squares (mean number of sperm heads examined ± SD: 212.12 ± 88.00). Variation in the sperm head shape (ASH) was treated as a binomial variable with heads classified either as normal (Fig. 1. A, D) or abnormal (Fig. 1. B, C, E, F). The proportion of ASH used for statistical analyses was estimated from 3 haemocytometer squares (mean number of sperm heads examined ± SD: 129.34 ± 52.01). All recorded phenotypic data are available in Supplementary Table S1.

**Fig. 1.**
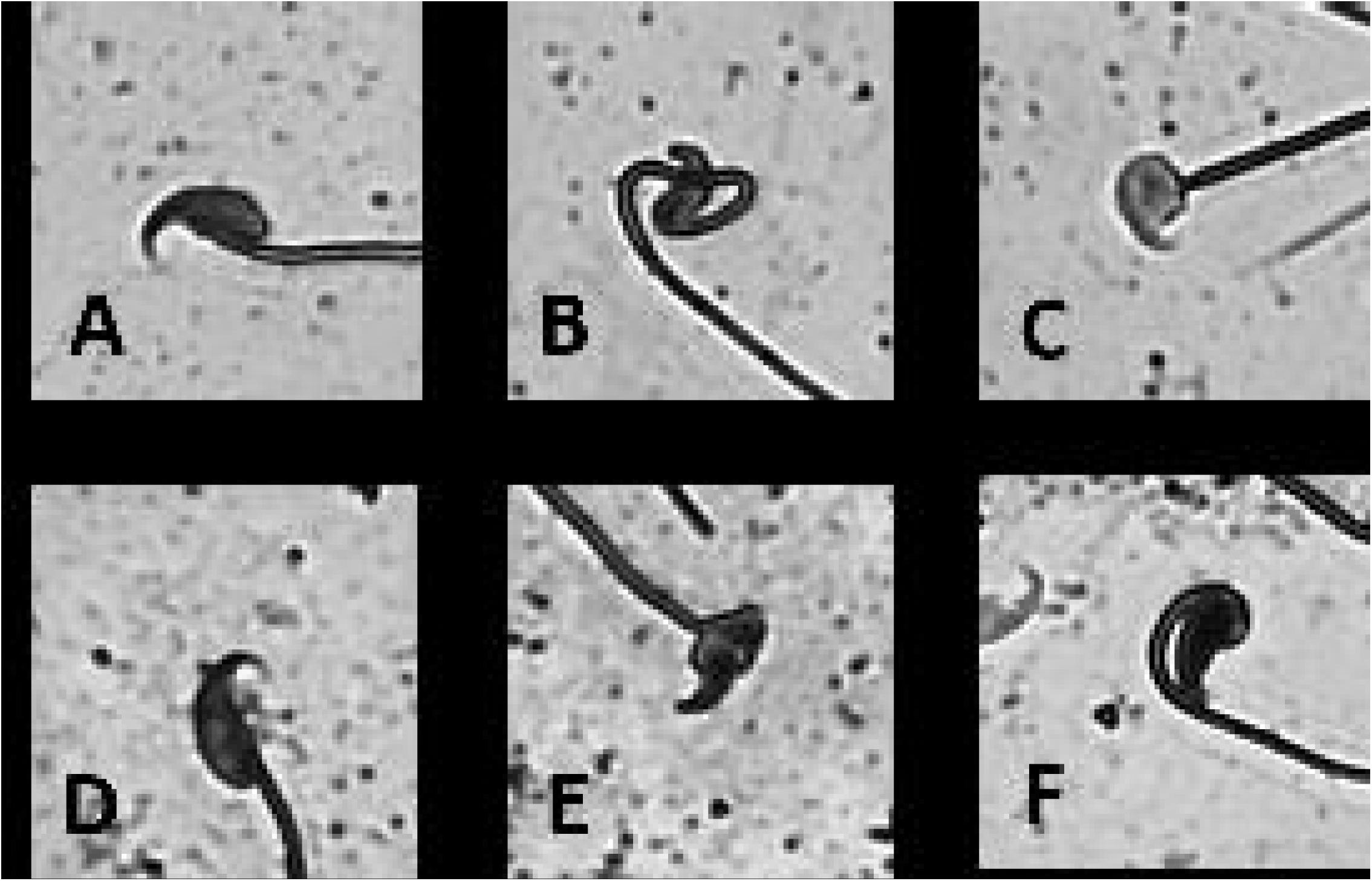
Classification of normal and abnormal sperm heads. Top panels show sperm heads of a (SCHUNT × MPB)F1 hybrid, lower panels heads of a (BIK × BULS)F1 hybrid. Figures A and D show normal heads, figures B, C and E, F show examples of abnormal sperm heads.

### Statistical analyses

Statistical analyses were performed in the R statistical environment (RStudio Team 2015, R Core Team 2018). The statistical models treat the sperm quality parameters (DSH and ASH) as dependent variables and cross type (*domesticus* × *domesticus, domesticus* × *musculus, musculus* × *domesticus, musculus* × *musculus*) as explanatory variables. Neither of DSH and ASH were distributed normally in F1 hybrids (Shapiro-Wilk normality test, P = 4.38×10^−10^ and P < 2.20×10^−16^, respectively). To improve the fit of DSH and ASH to normality we examined two transformations suggested in the literature for sperm head abnormalities: the arcsine square-root transformation (Kawai et al. 2006, White et al. 2011) and the Box-Cox power transformation utilized by Martincová et al. (2019).

Where data transformation achieved normality and variance homogeneity, parametric tests were used to analyse differentiation in sperm quality between the crosses. We applied linear modelling (function lm, from the R package *stats* (R Core Team 2018)) and a linear mixed-effects model fitted with the *lmer* function (R package lme4, Bates et al. (2015)). In mixed-effects models we asked whether unequal numbers of sires and dams used per individual strain (see Table 1 for exact numbers) disproportionally affected variance in the model and hence statistical inference. Both linear and mixed-effects models were tested for normality of residuals. Selection of the best model fit was based on the Akaike’s information criterion (ΔAIC) (Akaike 1974). For *post hoc* comparison of pairwise differences in between inter- and intrasubspecific crosses for the selected models we utilized the *glht* package (Bretz et al. 2010). Variables that did not conform to normal distribution were analysed using the non-parametric Kruskal-Walis test and *post hoc* Kruskal Neményi test for pair-wise comparisons of mean rank sums as implemented in the R package PMCMR (Pohlert 2014).

The proportion of explained variance associated with fixed effects (cross type) and with unequal contribution of sires and dams from the same strain was calculated using the approach described in Nakagawa & Schielzeth (2013). In addition, we estimated parameters for fixed effects and standard deviations for each random effect level. The parametric bootstrap was used to derive corresponding 95% confidence intervals.

## Results

The arcsine square-root transformation failed to achieve normality for both variables (Shapiro-Wilk normality test, P = 7.45×10^−5^ for DSH and P < 2.20×10^−16^ for ASH). Box-Cox power transformation achieved approximations to normality. For DSH, the optimal λ was estimated as 0.09. The transformed variable DSH.BC = (DSH^0.09^-1)/0.09 was in conformity with normality (Shapiro-Wilk normality test, P = 0.82) and displayed homogenous variances across the four crosses (Bartlett test, P = 0.42). The optimal λ for ASH was estimated at −0.50; however, the transformed variable ASH.BC = (ASH^−0.5^-1)/-0.5 displayed non-normal distribution (Shapiro-Wilk normality test, P = 0.00).

Frequencies of DSH within the four cross types are depicted in Fig 2 and detailed in Supplementary Table S2.1. Parametric model fitting revealed that the model with random effects of strain identity of sires and dams performed better than the linear model (ΔAIC = 6.29, P = 0.01; see Supplementary Table 2.2). Results of *post hoc* comparison of pairwise differences in frequencies of DSH in the mixed-effects model are summarized in Supplementary Table S2.3 and visualised with letters in Fig 2. Two patterns emerge from the results. The mean frequencies of DSH did not significantly differ between intrasubspecific crosses being respectively ∼10% and ∼12% in the *domesticus* and *musculus* F1 hybrids. However, the intersubspecific crosses displayed significant differences depending on cross type. The numbers of DSH were ∼8% in *domesticus* × *musculus* and 15% in *musculus* × *domesticus* males and the medians of DSH differ by more than 180% between them becoming highly significant (Tukey *post hoc* comparison, P <0.001).

**Fig. 2.**
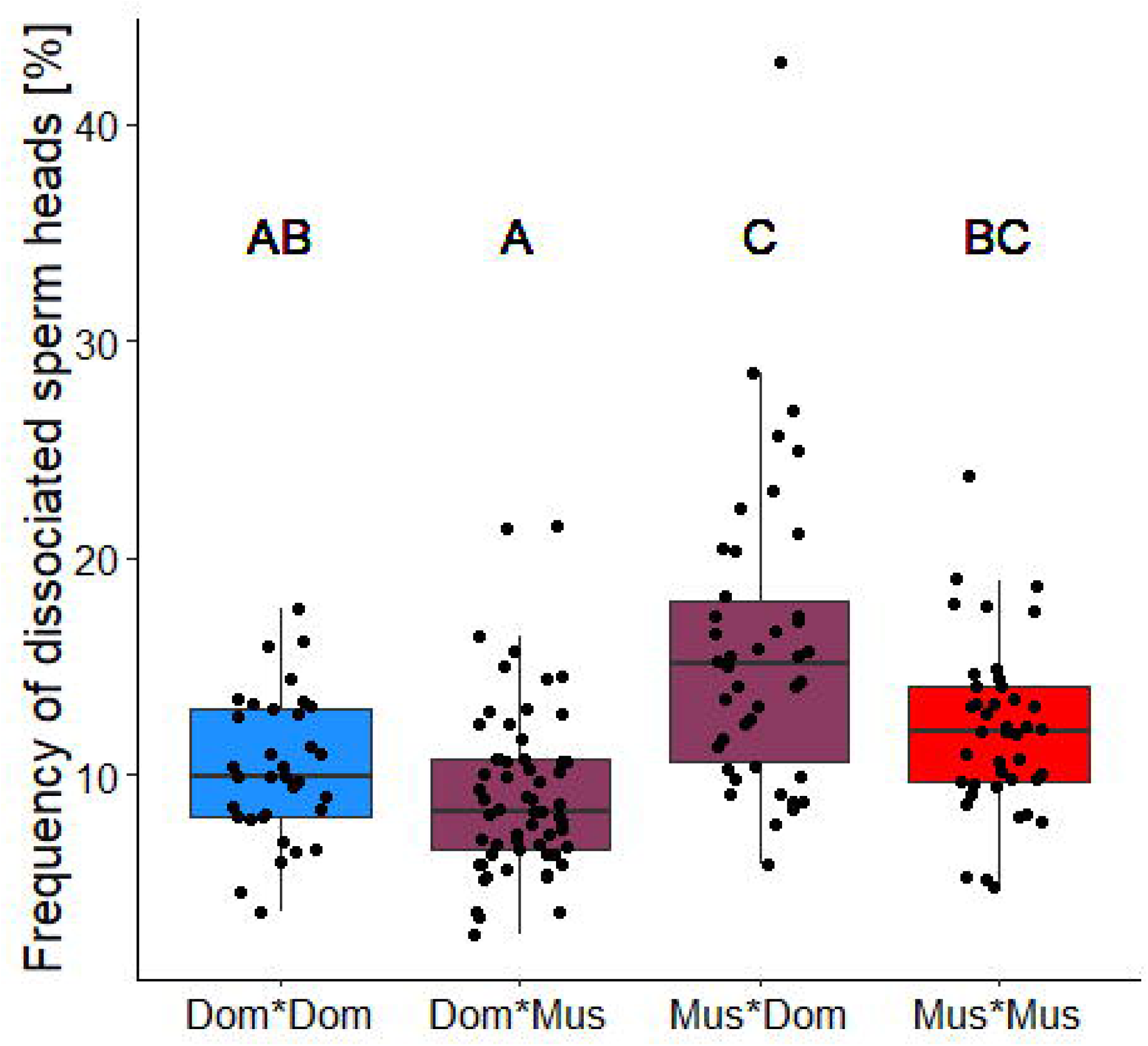
Frequency of dissociated sperm heads in intra- and intersubspecific crosses. Different letters indicate significant differences between crosses (see Supplementary Table 2.3). Medians, quartiles, 1.5 interquartile range and individual data are presented.

The explained fraction of variance attributed to cross type (fixed effects) in DSH was 23.41%, the fraction of variance attributed to strain origin of males was 12.35% and 8.77% to females and the proportion of unexplained variance was 55.46%.

Frequencies of sperm head abnormalities in F1 hybrids are shown in Fig. 3 and detailed in Supplementary Table S2.1. The nonparametric test detected cross type dependent differentiation (Kruskal-Wallis test, P = 4.95×10^−5^). The *post hoc* pairwise comparisons among cross types are in Supplementary Table S2.4 and shown with letters in Fig 3. The incidence of abnormal heads was almost the same in the intrasubspecific *musculus* and *domesticus* hybrids, reaching 6.98% and 7.10%, respectively. Hybridization increased ASH frequencies in both intersubspecific hybrids, though only the difference detected in the *domesticus* × *musculus* males was significant (ASH = 10.00%) and the *musculus* × *domesticus* males were intermediate (ASH = 8.13%). Males with severely deformed sperm head prevalence were rare: only 4 males (2.24%) displayed abnormal heads in frequencies higher than 50%, all of them originating from the *musculus* × *domesticus* cross.

**Fig. 3.**
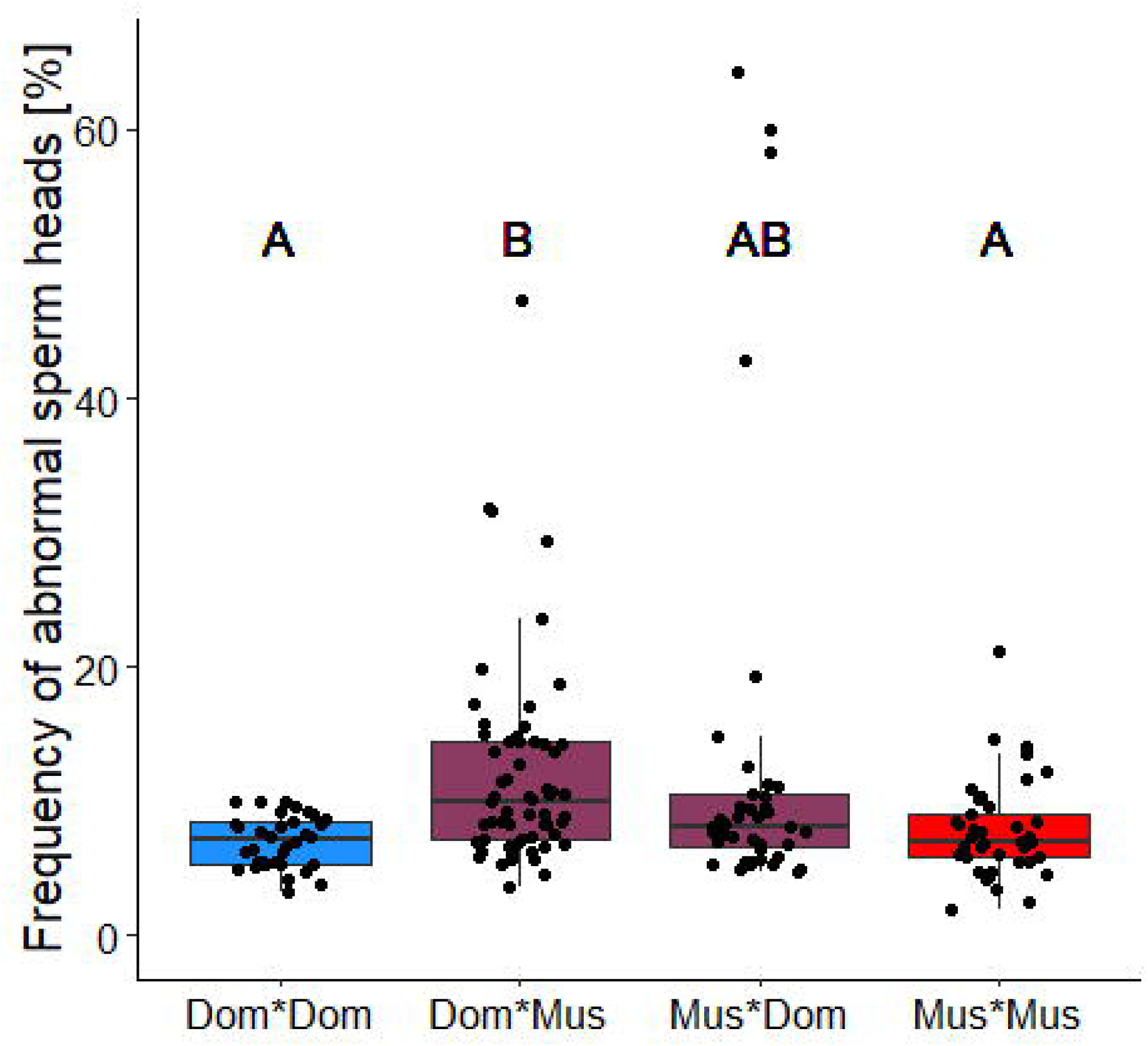
Frequency of abnormal sperm heads in intra- and intersubspecific crosses. Different letters indicate significant differences between crosses (see Supplementary Table 2.4). Medians, quartiles, 1.5 interquartile range and individual data are presented. One Mus*Dom male with ASH = 78.26% is out of the Y-axis limit.

## Discussion

Spermatogenesis is a tuned cascade of processes producing sperm and impairment of any phase of this process can affect fitness of males. The most severe impairments are caused by hybridization and in inter(sub)specific hybrids ultimately lead to sterility (Forejt et al. 2012). Using a wide spectrum of wild-derived strains, we investigated the effects of hybridization on sperm quality parameters that can potentially decrease fertilization success. We found that frequencies of abnormal and dissociated sperm heads do not differ between parental intrasubspecific hybrids (i.e. within the *musculus* and *domesticus* crosses). However, the proportions of DSH and ASH differ significantly between and are dependent on cross direction in intersubspecific hybrids.

We replicated the Martincová et al. (2019) results for subspecies-specific effects of hybridization on frequency of DSH, where 8 wild-derived strains showed significantly increased frequency of DSH in *musculus* × *domesticus* F1 hybrids in comparison with *domesticus* × *musculus* F1s. Our extension of DSH data to 29 strains strongly supports that this asymmetric effect is replicable among subspecific crosses. The *domesticus* × *musculus* F1s that display lower DSH frequency than the *musculus* × *musculus* males share their Y^musculus^ chromosome origin but differ in the subspecific origin of the X chromosome. Similarly, the *musculus* × *domesticus* F1s who have higher proportion of DSH than the *domesticus* × *domesticus* males share their Y^domesticus^ chromosome origin but possess different subspecific types of the X chromosome. This suggests that most of the DSH differences are associated with the X chromosome and modulated by interaction with the Y chromosome, which changes the direction of the response in the intersubspecific hybrids: the Y^domesticus^ increases and the Y^musculus^ decreases DSH frequency. Although the effects of autosomal loci cannot be tested in this experiment, the high variation observed in DSH data (the proportion of unexplained variance being over 55.46%) suggests the presence of polymorphic autosomal and sex-linked gene interactions.

Now considering ASH: frequencies were elevated in both reciprocal types of F1 hybrids in comparison with the parental F1 hybrids. We cannot replicate the 8 wild-derived strains results of Martincová et al. (2019), which suggested an increased frequency of ASH in the *musculus* × *domesticus* F1 hybrids. Those authors traced the causality of this variation to Y^domesticus^ from one locality interacting with autosomal or X-linked loci. None of these Y^domesticus^ chromosomes were however found in males displaying high rates of abnormal sperm (ASH > 30%, N = 8) in the current study. Moreover, among the four cross types the only significantly increased proportion of ASH was observed in the *domesticus* × *musculus* F1 hybrids, i.e., in the opposite direction of the selective advantage that would be required to explain the Y^musculus^ spread in the HMHZ. Hence, it seems that the incidence of ASH can be increased in intersubspecific mouse hybrids, but these effects appear polymorphic, with their occurrence conditioned by specific genotype(s).

For ASH genetic correlates between the sex chromosomes and sperm head abnormalities are well established (e.g. Ellis et al. 2005, Good et al. 2008, White et al. 2011, Cocquet et al. 2012, Campbell & Nachman 2014). However, these effect are not sex chromosome exclusive. In a laboratory congenic strain, B10.M, a high incidence of sperm-head morphological abnormalities was mapped to chromosomes 1 and 4 and no signal was detected on the X chromosome (Gotoh et al. 2012).

As far as we know, only one study has analysed genetic correlates for DSH between intersubspecific F1 hybrids (White et al., 2011). They found increased DSH incidence in *musculus* × *domesticus* F1s (represented respectively by wild-derived strains PWD and WSB) and this increase was more than 4 times higher in comparison with the reciprocal *domesticus* × *musculus* F1 hybrids. Quantitative trait loci associated with DSH were detected on proximal parts of chromosomes 15 (within a 16-48 Mb interval) and X (10-96 Mb interval) (White et al., 2011). The sex chromosome effects on DSH demonstrated in the current study apprear to be the strongests sperm trait effects found in experimental crosses using wild-derived mice (White et al., 2011, Martincová et al. 2019). Are these associations reflected by the physical positions of the genes participating in cohesion between the sperm head and tail?

The sperm head and tail are bridged by the myosin-based connecting piece. A comprehensive search detected various factors that interact with myosin subunits (Chen et al. 2016). The *Oaz3* gene that encodes ornithine decarboxylase antizyme 3 [located on chromosome 3], the *Odf1* gene encoding the outer dense fibre of sperm tails 1 [chromosome 15: 38.2 Mb] and the *Spata6* encoding spermatogenesis associated 6 [chromosome 4] participate in myosin-based microfilament formation (genomic positions were retrieved at http://www.informatics.jax.org/). Only SPATA6 forms a complex with myosin light and heavy chain subunits (e.g., MYL6 on chromosome 3) during connecting piece formation (Yuan et al. 2015). The *Oaz3* gene-encoded protein p12 targets phosphatase targeting subunit 3 Ppp1r16a (synonym MYPT3 on chromosome 15 [36.25 cM]) which is a regulator of activity of the protein phosphatases Ppp1cb (synonym PP1β on chromosome 5) and Ppp1cc (synonym PP1γ2 on chromosome 5) (Ruan et al. 2011). Linking of ODF1 to microtubules might occur via ODF1/SPAG5 [chromosome 11] / SPAG4 [chromosome 2] interaction. SPAG4 knock-out mice demonstrate that the SPAG4 itself is not essential for the formation of the sperm head-to-tail coupling apparatus, nevertheless, SPAG4 is required for tightening the sperm head-to-tail anchorage (Yang et al. 2018). Surprisingly, none of the genes noted above are located on the sex chromosomes, and only two autosomal genes (*Odf1* and Ppp1r16a) lie within the intervals identified for QTLs of DSH (White et al., 2011). This suggests that other sex chromosome-linked loci that can affect the tightness of sperm head-to-tail bonds remain to be determined.

The mechanism causing asymmetry in DSH frequencies between two types of intersubspecific hybrids might be considered in evolutionary context as a result of an arms race. The developmental environment of sperm in a male’s epididymis contains proteins encoded by genes on both the male’s X and Y chromosomes (though there are many more genes on the X chromosome). Any gene in the non-recombining X could increase its copy number in progeny of the male by selecting against Y-bearing sperm. One way to do so would be to encode a DSH promoter in the epididymis. A DSH-promoting X chromosome that also gives sperm carrying it (the same X) resistance to the promoter will win an X|Y arms race for passage through the epididymis. Y-bearing sperm will suffer disproportionate DSH until they gain ‘resistance’ to this X chromosome strategy – for example by increasing their own resistance to the DSH promoter. Such arms races are not uncommon in the house mouse (Cocquet et al. 2009, 2012, Ellis et al. 2011, Larson et al. 2017) and are, very likely, long-term features of the evolutionary process. Suppose, when the populations giving rise to *musculus* and *domesticus* were split (0.5 MYA), an on-going X|Y arms race in the epididymis was also split, following independent trajectories for half a million years before secondary contact (in both nature and laboratory crosses). It is highly unlikely that the arms races will come together in identical states. Coupled higher DHS promoter levels and resistance levels in *Mus* would be consistent with our observed results: Y^musculus^ sperm in a *domesticus* X epididymis combine high resistance with low promoter levels: low DSH. Y^domesticus^ sperm in a *musculus* X epididymis combine low resistance with high promoter levels: high DSH. This hypothesis has a testable prediction: DSH sperm should tend to be Y-bearing, not X, and a fluorescence *in-situ* hybridization protocol probing X and Y chromosomes can address this question. Finally, we note the hypothesis is consistent with both the Y^musculus^ invasion across the HMHZ and its associated distortion of the trapping sex ratio: ubiquitous low levels of X-induced DSH within each taxon would be consistent with female-biased sex ratios in ‘pure’ populations (Macholán et al. 2008). If the ‘Y^musculus^ sperm in a *domesticus* X epididymis combine high resistance with low promoter levels’ case eliminated DSH the prediction would be sex ratio parity – as observed in Y^musculus^ introgressed populations (Macholán et al. 2008).

A higher incidence of tailless sperm can also result as a consequence of sperm manipulation after its release from epididymis. For example, in patients in which semen analysis was normal, a minimal micromanipulation for ICSI resulted in decapitation of the spermatozoon during immobilization (Kamal et al. 1999). However, it would be difficult to explain the differences in DSH among the cross types we have observed, as all spermatozoa were manipulated using the same protocol.

Finally, by comparing two data transformations used in the literature we found that only the Box-Cox transformation was useful to approximate normality at least in some proportional data. We suggest this transformation as a preferable method for analysis of sperm quality parameters.

In conclusion, we found strong evidence that intersubspecific hybridization significantly affects sperm quality and these effects are dependent on sperm trait and cross type direction. The proportion of sperm head abnormalities was in general increased by hybridization, the frequency of dissociated sperm heads is increased in the *musculus* × *domesticus* F1 but decreased in the *domesticus* × *musculus* F1 hybrids. The reported differences in sperm quality between cross types may be highly relevant to male fitness in zones of secondary contact between the two house mouse subspecies. The uterus junction has been described as a barrier preventing deformed sperm reaching the eggs (Krzanowska, 1974, Nestor & Handel, 1984); consequently, hybrids with increased ASH may be targets of selection. On the other hand, head-tail dissociations have a very direct effect on fitness as tailless head cannot swim through the uterine environment to reach the female gametes. The cross direction asymmetry in frequency of dissociated sperm heads should favour the *M. m. musculus* Y chromosome in F1 male progeny. This is consistent with the spread of the *M. m. musculus* Y chromosome in nature across the hybrid zone between these two subspecies (Macholán et al. 2008, Albrechtová et al. 2012, Ďureje et al. 2012). The frequencies of both ASH and DSH observed in this study may appear low (medians ranging between 7.0-15.1%). However, their fitness consequences will be tested in nature in the presence of male-male sperm competition (Dean et al. 2006), and so have the potential to be very significant.

## Data availability

All data are available as electronic supplementary material

**Supplementary Material S1**. Phenotypic data for sperm traits in individual crosses. (.xlsx file)

**Supplementary Table 2.-2.4.** Descriptive statistics and tests including post hoc comparison for dissociated and abnormal sperm heads. (.docx file)

## Accession ID

House mouse (*Mus musculus*), the NCBI taxon ID 10090.

## Acknowledgements

We thank Jakub Kreisinger for statistical advice, and Lidka Rousková, Helena Hejlová, Iva Pospíšilová, and Jana Piálková for the welfare provided to the experimental mice.

This study was supported by the Czech Science Foundation (Projects 17-25320S and 19-12774S) and the ROZE program of the Czech Academy of Sciences. The funding agencies played no role in the design of the study, the collection, analysis, and interpretation of data and in writing the manuscript.

## References

Albrechtová J, Albrecht T, Baird SJE, Macholán M, Rudolfsen G, Munclinger P, Tucker PK, Piálek J (2012) Sperm-related phenotypes implicated in both maintenance and breakdown of a natural species barrier in the house mouse. Proc R Soc B-Biol Sci 279:4803–4810 https://doi.org/10.1098/rspb.2012.1802

Akaike H (1974) New look at statistical-model identification. IEEE Transact Automatic Control AC19:716–723 https://doi.org/10.1109/TAC.1974.1100705

Alund M., Immler S, Rice AM, Qvarnström A (2013) Low fertility of wild hybrid male flycatchers despite recent divergence. Biol Lett 9:20130169 https://doi.org/10.1098/rsbl.2013.0169

Baird SJE, Macholán M (2012) What can the *Mus musculus musculus/M. m. domesticus* hybrid zone tell us about speciation? In: Macholán M, Baird SJE, Munclinger P, Piálek J (eds.), Evolution of the House Mouse. Cambridge University Press, Cambridge:334–372

Bates D, Machler M, Bolker BM, Walker SC (2015) Fitting linear mixed-effects models using lme4. J Stat Software 67:1–48 https://doi.org/10.18637/jss.v067.i01

Bretz F, Hothorn T, Westfall P (2011) Multiple Comparisons Using R. CRC Press, Boca Raton

Campbell P, Nachman MW (2014) X-Y interactions underlie sperm head abnormality in hybrid male house mice. Genetics 196:1231–1240 https://doi.org/10.1534/genetics.114.161703

Chen SR, Batool A,. Wang YQ,. Hao XX, Chang CS, Cheng CY, Liu Y. X (2016) The control of male fertility by spermatid-specific factors: searching for contraceptive targets from spermatozoon’s head to tail. Cell Death & Dis 7:e2472 https://doi.org/10.1038/cddis.2016.344

Cocquet J, Ellis PJI, Yamauchi T, Mahadevaiah SK, Affara NA, Ward MA, Burgoyne PS (2009) The multicopy gene *Sly* represses the sex chromosomes in the male mouse germline after meiosis. PLoS Biol 7:e1000244 https://doi.org/10.1371/journal.pbio.1000244

Cocquet J, Ellis PJI, Mahadevaiah SK, Affara NA, Vaiman D, Burgoyne PS (2012) A genetic basis for a postmeiotic X versus Y chromosome intragenomic conflict in the mouse. PLoS Genetics 8:e1002900 https://doi.org/10.1371/journal.pgen.1002900

Coyne JA Orr HA (2004). Speciation. Sinauer Associates, Sunderland

Dean MD, Ardlie KG, Nachman MW (2006) The frequency of multiple paternity suggests that sperm competition is common in house mice (*Mus domesticus*). Mol Ecol 15:4141–4151 https://doi.org/10.1111/j.1365-294X.2006.03068.x

Ďureje L, Macholán M, Baird SJE, Piálek J (2012) The mouse hybrid zone in Central Europe: From morphology to molecules. Folia Zool 61:308–318

Ellis PJI, Clemente EJ, Ball P, Toure A, Ferguson L, Turner JMA, Loveland KL. Affara NA, Burgoyne PS (2005) Deletions on mouse Yq lead to upregulation of multiple X- and Y-linked transcripts in spermatids. Human Mol Genet 14:2705–2715 https://doi.org/10.1093/hmg/ddi304

Ellis PJI, Bacon J, Affara NA (2011) Association of *Sly* with sex-linked gene amplification during mouse evolution: a side effect of genomic conflict in spermatids? Human Mol Genet 20:3010–3021 https://doi.org/10.1093/hmg/ddr204

Forejt J, Piálek J, Trachtulec Z (2012) Hybrid male sterility genes in the mouse subspecific crosses. In: Evolution of the House Mouse, edited by. Macholán M, Baird SJE, Munclinger P, Piálek J, Cambridge: Cambridge Univ Press, p 482–503

Good JM, Dean MD, Nachman MW (2008) A complex genetic basis to X-linked hybrid male sterility between two species of house mice. Genetics 179: 2213–2228 https://doi.org/10.1534/genetics.107.085340

Gotoh H, Hirawatari K, Hanzawa N, Miura I, Wakana S (2012) QTL on mouse chromosomes 1 and 4 causing sperm-head morphological abnormality and male subfertility. Mamm Genome 23:399–403 https://doi.org/10.1007/s00335-012-9395-1

Gregorová S, Forejt J (2000) PWD/Ph and PWK/Ph inbred mouse strains of *Mus m. musculus* subspecies – a valuable resource of phenotypic variations and genomic polymorphisms. Folia Biol, Praha 46:31–41

Kamal A, Mansour R, Fahmy I, Serour G, Rhodes C, Aboulghar M (1999) Easily decapitated spermatozoa defect: a possible cause of unexplained infertility. Human Reprod 14:2791–2795 https://doi.org/10.1093/humrep/14.11.2791

Kawai Y, Hata T, Suzuki O, Matsuda J (2006) The relationship between sperm morphology and in vitro fertilization ability in mice. J Reprod Dev 52:561–568 https://doi.org/10.1262/jrd.18023

Krzanowska H (1974) The passage of abnormal spermatozoa through the uterotubal junction of the mouse. J Reprod Fert 34:81–90

Larson EL, Keeble S, Vanderpool D, Dean MD, Good JM (2017) The composite regulatory basis of the large X-effect in mouse speciation. Mol Biol Evol 34:282–295 https://doi.org/10.1093/molbev/msw243

Macholán M, Munclinger P, Šugerková M, Dufková P, Bímová B, Božíková E, Zima J, Piálek J (2007) Genetic analysis of autosomal and X-linked markers across a mouse hybrid zone. Evolution 61:746–771 https://doi.org/10.1111/j.1558-5646.2007.00065.x

Macholán M, Baird SJE, Munclinger P, Dufková P, Bímová B, Piálek J (2008) Genetic conflict outweighs heterogametic incompatibility in the mouse hybrid zone? BMC Evol Biol 8:271 https://doi.org/10.1186/1471-2148-8-271

Martincová I., Ďureje L., Kreisinger J., Macholán M., Piálek J. 2019. Phenotypic effects of the Y chromosome are variable and structured in hybrids among house mouse recombinant lines. Ecol Evol: 9:6124–6137. https://doi.org/10.1002/ece3.5196

Nakagawa S, Schielzeth H (2013) A general and simple method for obtaining *R*^*2*^ from generalized linear mixed-effects models. Methods Ecol Evol 4:133–142 https://doi.org/10.1111/j.2041-210x.2012.00261.x

Nestor A, Handel, MA (1984) The transport of morphologically abnormal sperm in the female reproductive tract of mice. Gamete Res 10:119–125 https://doi.org/10.1002/mrd.1120100204

Piálek J, Vyskočilová M, Bímová B, Havelková D, Piálková J, Dufková P, Bencová V, Ďureje L, Albrecht T, Hauffe HC, Macholán M, Munclinger P, Storchová R, Zajícová A, Holáň V, Gregorová S, Forejt J (2008) Development of unique house mouse resources suitable for evolutionary studies of speciation. J Heredity 99:34–44 https://doi.org/10.1093/jhered/esm083

Pohlert T (2014) The pairwise multiple comparison of mean ranks package (PMCMR). R package. https://CRAN.R-project.org/package=PMCMR

R Core Team (2018). R: A language and environment for statistical computing. R Foundation for Statistical Computing, Vienna, Austria. URL: https://www.R-project.org/

RStudio Team (2015) RStudio: Integrated Development for R. RStudio, Inc., Boston. Retrieved from http://www.rstudio.com/

Ruan Y, Cheng M, Ou Y, Oko R, van der Hoorn FA (2011) Ornithine decarboxylase antizyme *Oaz3* modulates protein phosphatase activity. J Biol Chem 286:29417–29427

Vyskočilová M, Trachtulec Z, Forejt J, Piálek J (2005) Does geography matter in hybrid sterility in house mice? Biol J Linn Soc 84:663–674 https://doi.org/10.1111/j.1095-8312.2005.00463.x

White M, Steffy B, Wiltshire T, Payseur, BA (2011) Genetic dissection of a key reproductive barrier between nascent species of house mice. Genetics 189:289–304 https://doi.org/10.1534/genetics.111.129171

Yuan S, Stratton CJ, Bao J, Zheng H, Bhetwal BP, Yanagimachi R, Yan W (2015) *Spata6* is required for normal assembly of the sperm connecting piece and tight head-tail conjunction. Proc Natl Acad Sci USA 112:E430–E439 https://doi.org/10.1073/pnas.1424648112

Yang K, Adham IM, Meinhardt A, Hoyer-Fender S (2018) Ultra-structure of the sperm head-to-tail linkage complex in the absence of the spermatid-specific LINC component SPAG4. Histochemistry Cell Biol 150:49–59 https://doi.org/10.1007/s00418-018-1668-7

