## Supplementary Material S1 for "Sperm quality parameters are increased and asymmetric in house mouse hybrids"

| Cross | Dam | Sir | Birth | PIN | Sex | Age | BW | N Sperm | N Dissoc | N Norm |
|---|---|---|---|---|---|---|---|---|---|---|
| DyM*Dom | BL6 | DDO | 03/12/2018 | EX2239 | M | 60 | 25.42 | 314 | 20 | 294 |
| DyM*Dom | BL6 | DGA | 26/11/2018 | EX2172 | M | 60 | 23.49 | 361 | 21 | 340 |
| DyM*Dom | BL6 | DJO | 21/11/2018 | EX2147 | M | 60 | 23.62 | 290 | 38 | 252 |
| DyM*Dom | BL6 | STAIL | 06/06/2017 | SY2475 | M | 60 | 25.65 | 329 | 33 | 296 |
| DyM*Dom | BL6 | STRA | 08/03/2016 | ISII6378 | M | 60 | 23.81 | 311 | 33 | 278 |
| DyM*Dom | BL6 | STRB | 28/11/2016 | ISII9900 | M | 60 | 22.36 | 316 | 47 | 269 |
| DyM*Mus | BL6 | PWD | 07/10/2018 | EX1903 | M | 60 | 22.43 | 6 | 2 | 4 |
| DyM*Mus | BL6 | SKE | 27/06/2018 | EX1313 | M | 60 | 20.24 | 130 | 10 | 120 |
| DyM*Mus | BL6 | STUF | 27/12/2018 | EX2351 | M | 60 | 25.46 | 237 | 12 | 225 |
| DyM*Mus | BL6 | STUS | 15/09/2018 | EX1763 | M | 60 | 20.08 | 0 | 0 | 0 |
| DyM*DyM | SPLY | BL6 | 26/02/2017 | SY1040 | M | 60 | 27.32 | 374 | 26 | 348 |
| DyM*DyM | SPLY | SPLY | 10/05/2016 | ISII7257 | M | 60 | 20.65 | 201 | 38 | 163 |
| DyM*Dom | SPLY | DCP | 07/02/2018 | EX554 | M | 60 | 22.16 | 281 | 29 | 252 |
| DyM*Dom | SPLY | SCHUNT | 05/10/2016 | ISII9194 | M | 60 | 19.49 | 232 | 33 | 199 |
| DyM*Dom | SPLY | SIN | 13/10/2017 | EX174 | M | 60 | 21.9 | 308 | 16 | 292 |
| DyM*Dom | SPLY | SIT | 22/01/2018 | EX497 | M | 60 | 18.99 | 256 | 20 | 236 |
| DyM*Dom | SPLY | STAIL | 11/01/2017 | SY509 | M | 60 | 24.07 | 257 | 32 | 225 |
| DyM*Dom | SPLY | STRA | 29/12/2016 | SY375 | M | 60 | 23.23 | 188 | 17 | 171 |
| DyM*Dom | SPLY | STRB | 12/12/2016 | SY102 | M | 60 | 17.4 | 184 | 23 | 161 |
| DyM*Mus | SPLY | BUSNA | 05/03/2018 | EX657 | M | 61 | 17.66 | 199 | 20 | 179 |
| DyM*Mus | SPLY | MPB | 29/08/2018 | EX1694 | M | 60 | 20.46 | 105 | 5 | 100 |
| DyM*Mus | SPLY | PWD | 31/08/2016 | ISII8777 | M | 60 | 20.16 | 329 | 39 | 290 |
| DyM*Mus | SPLY | STUF | 30/05/2017 | SY2344 | M | 60 | 22.84 | 205 | 19 | 186 |
| DyM*Mus | SPLY | STUS | 25/02/2018 | EX626 | M | 60 | 19.4 | 89 | 14 | 75 |
| Dom*Dom | BIK | DCP | 19/04/2018 | EX863 | M | 60 | 18.95 | 229 | 30 | 199 |
| Dom*Mus | BIK | BULS | 12/01/2019 | EX2444 | M | 60 | 19.9 | 135 | 5 | 130 |
| Dom*Mus | BIK | MPB | 06/07/2018 | EX1359 | M | 60 | 16.77 | 63 | 4 | 59 |
| Dom*Mus | BIK | PWD | 22/09/2018 | EX1794 | M | 60 | 18.62 | 185 | 18 | 167 |
| Dom*Mus | BIK | STUF | 04/10/2018 | EX1883 | M | 60 | 19.82 | 136 | 9 | 127 |
| Dom*Mus | BIK | STUS | 08/08/2018 | EX1564 | M | 60 | 19.97 | 86 | 10 | 76 |
| Dom*Dom | BZO | SIT | 10/08/2018 | EX1596 | M | 60 | 19.11 | 220 | 8 | 212 |
| Dom*Mus | BZO | BULS | 02/07/2018 | EX1347 | M | 60 | 15.74 | 121 | 13 | 108 |
| Dom*Mus | BZO | PWD | 18/12/2018 | EX2331 | M | 60 | 16.73 | 189 | 7 | 182 |
| Dom*Mus | BZO | STUF | 03/11/2018 | EX2069 | M | 60 | 17.89 | 219 | 16 | 203 |
| Dom*DyM | DCP | BL6 | 14/12/2018 | EX2313 | M | 60 | 23.77 | 246 | 15 | 231 |
| Dom*Dom | DCP | BIK | 01/07/2018 | EX1335 | M | 60 | 15.61 | 166 | 10 | 156 |
| Dom*Dom | DCP | DCP | 22/01/2018 | SY8995 | M | 60 | 15.93 | 163 | 26 | 137 |
| Dom*Mus | DCP | BULS | 14/04/2018 | EX828 | M | 60 | 21.29 | 208 | 11 | 197 |
| Dom*Mus | DCP | BUSNA | 01/11/2018 | EX2059 | M | 60 | 18.29 | 192 | 13 | 179 |
| Dom*Mus | DCP | MHB | 13/09/2018 | EX1759 | M | 60 | 25.41 | 254 | 16 | 238 |
| Dom*Mus | DCP | PWD | 06/08/2018 | EX1553 | M | 60 | 19.03 | 222 | 15 | 207 |
| Dom*DyM | DDO | BL6 | 01/01/2019 | EX2375 | M | 60 | 26.21 | 207 | 14 | 193 |
| Dom*Dom | DDO | WLA | 07/11/2018 | EX2086 | M | 60 | 19.51 | 220 | 18 | 202 |

| | | | | | | | | | | |
|---|---|---|---|---|---|---|---|---|---|---|
| Dom*Mus | DDO | PWD | 28/08/2018 | EX1688 | M | 60 | 20.86 | 119 | 9 | 110 |
| Dom*Mus | DDO | SKA | 13/12/2018 | EX2304 | M | 60 | 16.54 | 38 | 1 | 37 |
| Dom*Mus | DDO | STUS | 22/06/2018 | EX1286 | M | 60 | 20.51 | 31 | 4 | 27 |
| Dom*DyM | DGA | BL6 | 23/10/2018 | EX1972 | M | 60 | 23.69 | 282 | 18 | 264 |
| Dom*Dom | DGA | SIT | 29/07/2018 | EX1502 | M | 60 | 17.06 | 288 | 30 | 258 |
| Dom*Mus | DGA | BUSNA | 09/12/2018 | EX2279 | M | 60 | 15.83 | 251 | 13 | 238 |
| Dom*Mus | DGA | MPB | 08/09/2018 | EX1747 | M | 60 | 16.26 | 266 | 24 | 242 |
| Dom*Mus | DGA | PWD | 30/11/2018 | EX2222 | M | 60 | 16.08 | 193 | 13 | 180 |
| Dom*Mus | DGA | SKE | 02/01/2019 | EX2383 | M | 60 | 16.84 | 168 | 14 | 154 |
| Dom*Mus | DGA | STUS | 17/11/2018 | EX2135 | M | 60 | 17.7 | 239 | 14 | 225 |
| Dom*Dom | DJO | DJO | 08/01/2019 | SX7832 | M | 60 | 14.25 | 93 | 15 | 78 |
| Dom*Mus | DJO | BUSNA | 16/01/2019 | EX2462 | M | 60 | 19.17 | 243 | 21 | 222 |
| Dom*Mus | DJO | STUF | 26/10/2018 | EX2006 | M | 60 | 19.91 | 208 | 21 | 187 |
| Dom*DyM | SCHUNT | BL6 | 11/03/2016 | ISII6401 | M | 60 | 24.85 | 347 | 31 | 316 |
| Dom*DyM | SCHUNT | SPLY | 03/04/2016 | ISII6887 | M | 60 | 19.95 | 415 | 49 | 366 |
| Dom*Dom | SCHUNT | BIK | 01/10/2018 | EX1848 | M | 60 | 18.08 | 310 | 34 | 276 |
| Dom*Dom | SCHUNT | DDO | 08/09/2018 | EX1748 | M | 60 | 23.83 | 346 | 24 | 322 |
| Dom*Dom | SCHUNT | DIK | 17/10/2018 | EX1949 | M | 60 | 21.03 | 303 | 30 | 273 |
| Dom*Dom | SCHUNT | SCHUNT | 20/03/2016 | ISII6499 | M | 60 | 20.95 | 204 | 27 | 177 |
| Dom*Dom | SCHUNT | SIN | 09/01/2018 | EX455 | M | 60 | 21.43 | 332 | 28 | 304 |
| Dom*Dom | SCHUNT | SIT | 26/05/2018 | EX1106 | M | 60 | 22.42 | 273 | 27 | 246 |
| Dom*Dom | SCHUNT | STRA | 25/10/2016 | ISII9412 | M | 60 | 19.5 | 187 | 27 | 160 |
| Dom*Dom | SCHUNT | STRB | 02/04/2017 | SY1508 | M | 60 | 21.79 | 363 | 29 | 334 |
| Dom*Mus | SCHUNT | BULS | 12/05/2017 | SY2112 | M | 61 | 20.07 | 214 | 35 | 179 |
| Dom*Mus | SCHUNT | MCZ | 10/01/2019 | EX2416 | M | 60 | 20.52 | 28 | 6 | 22 |
| Dom*Mus | SCHUNT | MPB | 21/01/2019 | EX2475 | M | 60 | 18.9 | 97 | 10 | 87 |
| Dom*Mus | SCHUNT | PWD | 29/09/2017 | EX124 | M | 60 | 17.1 | 267 | 39 | 228 |
| Dom*Mus | SCHUNT | SENK | 02/02/2018 | EX582 | M | 60 | 18.01 | 217 | 23 | 194 |
| Dom*Mus | SCHUNT | SKA | 22/02/2018 | EX615 | M | 60 | 18.21 | 199 | 30 | 169 |
| Dom*Mus | SCHUNT | SKE | 18/12/2018 | EX2329 | M | 60 | 16.55 | 147 | 8 | 139 |
| Dom*Mus | SCHUNT | STUF | 23/06/2016 | ISII7886 | M | 60 | 18.83 | 301 | 32 | 269 |
| Dom*Mus | SCHUNT | STUS | 17/02/2017 | SY900 | M | 60 | 17.26 | 166 | 26 | 140 |
| Dom*Dom | SIN | BZO | 03/08/2018 | EX1529 | M | 60 | 17.35 | 236 | 11 | 225 |
| Dom*Mus | SIN | MPB | 01/09/2018 | EX1716 | M | 61 | 17.79 | 145 | 5 | 140 |
| Dom*Mus | SIN | PWD | 08/11/2017 | EX294 | M | 60 | 17.02 | 226 | 16 | 210 |
| Dom*Mus | SIN | STUF | 26/02/2018 | EX633 | M | 60 | 17.66 | 222 | 14 | 208 |
| Dom*DyM | SIT | BL6 | 22/11/2018 | EX2163 | M | 60 | 25.37 | 302 | 28 | 274 |
| Dom*Dom | SIT | BIK | 30/06/2018 | EX1332 | M | 60 | 19.04 | 226 | 29 | 197 |
| Dom*Dom | SIT | DCP | 23/03/2018 | EX719 | M | 60 | 18.14 | 173 | 14 | 159 |
| Dom*Dom | SIT | SCHUNT | 07/07/2018 | EX1378 | M | 60 | 18.74 | 348 | 33 | 315 |
| Dom*Dom | SIT | STRA | 30/06/2018 | EX1331 | M | 60 | 26.17 | 268 | 28 | 240 |
| Dom*Dom | SIT | STRB | 01/12/2018 | EX2223 | M | 60 | 22.56 | 334 | 27 | 307 |
| Dom*Dom | SIT | WLA | 30/07/2018 | EX1512 | M | 60 | 17.65 | 279 | 25 | 254 |
| Dom*Mus | SIT | BULS | 10/04/2018 | EX811 | M | 60 | 17.62 | 112 | 10 | 102 |
| Dom*Mus | SIT | BUSNA | 07/10/2018 | EX1902 | M | 60 | 16.92 | 179 | 15 | 164 |
| Dom*Mus | SIT | MGA | 17/11/2018 | EX2136 | M | 60 | 23.12 | 254 | 18 | 236 |
| Dom*Mus | SIT | MPB | 25/09/2018 | EX1809 | M | 60 | 20.33 | 142 | 11 | 131 |
| Dom*Mus | SIT | SKA | 04/01/2018 | EX436 | M | 60 | 22.46 | 146 | 18 | 128 |
| Dom*Mus | SIT | STUS | 10/06/2018 | EX1203 | M | 60 | 16.93 | 79 | 17 | 62 |
| Dom*DyM | STAIL | BL6 | 08/07/2017 | SY3051 | M | 60 | 26.28 | 366 | 45 | 321 |

| | | | | | | | | | | |
|---|---|---|---|---|---|---|---|---|---|---|
| Dom*Dom | STAIL | STAIL | 23/08/2017 | SY4537 | M | 60 | 13.83 | 238 | 32 | 206 |
| Dom*Mus | STAIL | BULS | 30/06/2017 | SY2944 | M | 60 | 20.59 | 141 | 18 | 123 |
| Dom*Mus | STAIL | MGA | 28/11/2018 | EX2206 | M | 60 | 19.15 | 56 | 6 | 50 |
| Dom*Mus | STAIL | SENK | 12/10/2017 | EX173 | M | 60 | 18.59 | 62 | 5 | 57 |
| Dom*Mus | STAIL | STUF | 03/11/2017 | EX273 | M | 60 | 22.79 | 198 | 21 | 177 |
| Dom*DyM | STRA | SPLY | 31/10/2016 | ISII9491 | M | 60 | 25.14 | 373 | 44 | 329 |
| Dom*Dom | STRA | BZO | 09/10/2018 | EX1914 | M | 60 | 26.32 | 291 | 25 | 266 |
| Dom*Dom | STRA | DCP | 11/07/2018 | EX1415 | M | 60 | 24.35 | 330 | 32 | 298 |
| Dom*Dom | STRA | DGA | 12/11/2018 | EX2104 | M | 60 | 23.01 | 353 | 23 | 330 |
| Dom*Dom | STRA | SCHUNT | 18/04/2017 | SY1789 | M | 60 | 24.16 | 346 | 38 | 308 |
| Dom*Dom | STRA | SIT | 14/05/2018 | EX1024 | M | 60 | 24.15 | 344 | 39 | 305 |
| Dom*Dom | STRA | STRA | 15/02/2016 | ISII6044 | M | 60 | 25.4 | 221 | 30 | 191 |
| Dom*Mus | STRA | BULS | 15/11/2018 | EX2124 | M | 60 | 20.27 | 175 | 12 | 163 |
| Dom*Mus | STRA | BUSNA | 08/11/2013 | DC62 | M | 60 | 18.5 | 168 | 17 | 151 |
| Dom*Mus | STRA | PWD | 23/02/2018 | EX619 | M | 60 | 24.05 | 303 | 27 | 276 |
| Dom*Mus | STRA | SENK | 06/05/2017 | SY2044 | M | 60 | 23.1 | 221 | 16 | 205 |
| Dom*Mus | STRA | SKA | 13/12/2018 | EX2305 | M | 60 | 22.2 | 158 | 13 | 145 |
| Dom*Mus | STRA | SKE | 10/10/2018 | EX1918 | M | 60 | 17.56 | 168 | 22 | 146 |
| Dom*Mus | STRA | STUF | 22/04/2017 | SY1847 | M | 60 | 27.99 | 252 | 27 | 225 |
| Dom*Mus | STRA | STUS | 12/10/2016 | ISII9260 | M | 60 | 21.44 | 214 | 31 | 183 |
| Dom*DyM | STRB | BL6 | 03/05/2017 | SY2021 | M | 60 | 30.92 | 354 | 39 | 315 |
| Dom*DyM | STRB | SPLY | 22/02/2016 | ISII6131 | M | 60 | 19.24 | 320 | 25 | 295 |
| Dom*Dom | STRB | DCP | 12/06/2018 | EX1206 | M | 60 | 21.47 | 366 | 24 | 342 |
| Dom*Dom | STRB | SCHUNT | 08/12/2016 | SY43 | M | 60 | 20.03 | 282 | 50 | 232 |
| Dom*Dom | STRB | STRA | 15/12/2016 | SY139 | M | 60 | 23.56 | 318 | 42 | 276 |
| Dom*Dom | STRB | STRB | 19/05/2016 | ISII7375 | M | 60 | 20.62 | 196 | 25 | 171 |
| Dom*Mus | STRB | BULS | 05/03/2014 | DC100 | M | 60 | 19.79 | 169 | 10 | 159 |
| Dom*Mus | STRB | MPB | 25/12/2018 | EX2349 | M | 60 | 18.08 | 248 | 14 | 234 |
| Dom*Mus | STRB | PWD | 30/01/2017 | SY687 | M | 60 | 22.15 | 264 | 14 | 250 |
| Dom*Mus | STRB | SENK | 13/11/2017 | EX306 | M | 60 | 19.04 | 257 | 24 | 233 |
| Dom*Mus | STRB | STUF | 19/05/2017 | SY2214 | M | 62 | 23.34 | 294 | 23 | 271 |
| Dom*Mus | STRB | STUS | 25/11/2013 | DC67 | M | 60 | 19.5 | 97 | 12 | 85 |
| Dom*Dom | WLA | WLA | 11/09/2018 | SX5161 | M | 60 | 14.3 | 261 | 26 | 235 |
| Dom*Mus | WLA | BULS | 29/11/2018 | EX2212 | M | 60 | 13.84 | 170 | 10 | 160 |
| Dom*Mus | WLA | BUSNA | 24/08/2018 | EX1676 | M | 60 | 14.94 | 228 | 19 | 209 |
| Dom*Mus | WLA | STUF | 17/04/2018 | EX857 | M | 60 | 17.95 | 273 | 27 | 246 |
| Mus*DyM | BULS | SPLY | 08/08/2018 | EX1565 | M | 60 | 21.15 | 117 | 15 | 102 |
| Mus*Dom | BULS | BIK | 28/05/2018 | EX1124 | M | 60 | 17.42 | 139 | 23 | 116 |
| Mus*Dom | BULS | DCP | 12/03/2018 | EX674 | M | 60 | 17.75 | 185 | 26 | 159 |
| Mus*Dom | BULS | SCHUNT | 18/03/2017 | SY1339 | M | 60 | 19.05 | 263 | 45 | 218 |
| Mus*Dom | BULS | SIN | 09/07/2018 | EX1392 | M | 60 | 17.26 | 12 | 3 | 9 |
| Mus*Dom | BULS | SIT | 01/02/2018 | EX543 | M | 60 | 18.26 | 7 | 3 | 4 |
| Mus*Dom | BULS | STRA | 03/04/2017 | SY1530 | M | 60 | 16.95 | 207 | 42 | 165 |
| Mus*Dom | BULS | STRB | 31/10/2016 | ISII9493 | M | 60 | 14.94 | 193 | 43 | 150 |
| Mus*Mus | BULS | BULS | 19/03/2016 | ISII6489 | M | 60 | 15.69 | 108 | 13 | 95 |
| Mus*Mus | BULS | BUSNA | 01/02/2018 | EX542 | M | 60 | 13.62 | 159 | 13 | 146 |
| Mus*Mus | BULS | MAM | 23/12/2018 | EX2341 | M | 60 | 16.01 | 170 | 9 | 161 |
| Mus*Mus | BULS | PWD | 13/06/2017 | SY2585 | M | 60 | 16.97 | 164 | 20 | 144 |
| Mus*Mus | BULS | SKA | 25/03/2018 | EX740 | M | 60 | 15.69 | 205 | 36 | 169 |
| Mus*Mus | BULS | SKE | 13/10/2018 | EX1931 | M | 60 | 17.61 | 136 | 7 | 129 |

| | | | | | | | | | | |
|---|---|---|---|---|---|---|---|---|---|---|
| Mus*Mus | BULS | STUS | 19/01/2014 | DC86 | M | 60 | 17.05 | 142 | 27 | 115 |
| Mus*Dom | BUSNA | DGA | 27/12/2018 | EX2359 | M | 60 | 18.42 | 355 | 31 | 324 |
| Mus*Dom | BUSNA | DJO | 20/09/2018 | EX1779 | M | 60 | 21.9 | 104 | 24 | 80 |
| Mus*Dom | BUSNA | SCHUNT | 08/02/2018 | EX555 | M | 60 | 22.07 | 339 | 46 | 293 |
| Mus*Dom | BUSNA | STRA | 14/03/2017 | SY1252 | M | 60 | 21.28 | 263 | 48 | 215 |
| Mus*Dom | BUSNA | STRB | 04/09/2016 | ISII8810 | M | 60 | 21.75 | 256 | 39 | 217 |
| Mus*Mus | BUSNA | BUSNA | 27/03/2016 | ISII6663 | M | 60 | 15.78 | 252 | 34 | 218 |
| Mus*Mus | BUSNA | MAM | 03/01/2019 | EX2392 | M | 60 | 19.11 | 309 | 25 | 284 |
| Mus*Mus | BUSNA | PWD | 26/03/2017 | SY1430 | M | 60 | 19.15 | 242 | 31 | 211 |
| Mus*Mus | BUSNA | SKE | 12/12/2018 | EX2290 | M | 60 | 15.82 | 221 | 21 | 200 |
| Mus*Mus | BUSNA | STUF | 03/11/2013 | DC50 | M | 60 | 19.78 | 185 | 26 | 159 |
| Mus*DyM | MAM | BL6 | 29/12/2018 | EX2362 | M | 60 | 24.43 | 253 | 21 | 232 |
| Mus*Dom | MAM | SIT | 21/09/2018 | EX1791 | M | 60 | 22.65 | 245 | 24 | 221 |
| Mus*DyM | MCZ | BL6 | 18/01/2019 | EX2469 | M | 60 | 20.88 | 133 | 20 | 113 |
| Mus*Dom | MGA | STRA | 07/11/2018 | EX2087 | M | 60 | 22.74 | 266 | 40 | 226 |
| Mus*Mus | MGA | PWD | 06/12/2018 | EX2270 | M | 60 | 15.11 | 203 | 20 | 183 |
| Mus*Mus | MGA | STUS | 26/11/2018 | EX2173 | M | 60 | 16.62 | 240 | 23 | 217 |
| Mus*Dom | MHB | DDO | 23/08/2018 | EX1669 | M | 60 | 23.1 | 320 | 28 | 292 |
| Mus*DyM | MPB | BL6 | 13/11/2018 | EX2113 | M | 60 | 20.58 | 127 | 17 | 110 |
| Mus*Dom | MPB | BIK | 20/12/2018 | EX2337 | M | 60 | 17.14 | 231 | 23 | 208 |
| Mus*Dom | MPB | DGA | 21/09/2018 | EX1792 | M | 60 | 17.15 | 285 | 40 | 245 |
| Mus*Dom | MPB | SCHUNT | 18/09/2018 | EX1774 | M | 60 | 18.35 | 172 | 27 | 145 |
| Mus*Dom | MPB | SIN | 15/07/2018 | EX1447 | M | 60 | 17.67 | 168 | 26 | 142 |
| Mus*Dom | MPB | STRA | 11/09/2018 | EX1751 | M | 60 | 18.02 | 127 | 16 | 111 |
| Mus*Mus | MPB | BUSNA | 23/08/2018 | EX1667 | M | 60 | 17 | 284 | 31 | 253 |
| Mus*Mus | MPB | PWD | 12/08/2018 | EX1609 | M | 60 | 15.81 | 124 | 6 | 118 |
| Mus*Mus | MPB | SKE | 10/09/2018 | EX1749 | M | 60 | 12.84 | 196 | 18 | 178 |
| Mus*DyM | PWD | BL6 | 19/01/2017 | SY568 | M | 60 | 23.75 | 0 | 0 | 0 |
| Mus*DyM | PWD | SPLY | 20/06/2017 | SY2754 | M | 60 | 22.71 | 165 | 29 | 136 |
| Mus*Dom | PWD | BIK | 14/12/2018 | EX2312 | M | 60 | 18.12 | 232 | 27 | 205 |
| Mus*Dom | PWD | DCA | 28/12/2018 | EX2361 | M | 60 | 17.11 | 226 | 28 | 198 |
| Mus*Dom | PWD | DJO | 23/12/2018 | EX2342 | M | 60 | 15.48 | 14 | 4 | 10 |
| Mus*Dom | PWD | DOT | 18/12/2018 | EX2330 | M | 60 | 22.33 | 233 | 36 | 197 |
| Mus*Dom | PWD | SCHUNT | 22/05/2016 | ISII7413 | M | 60 | 18.75 | 234 | 48 | 186 |
| Mus*Dom | PWD | STAIL | 14/11/2016 | ISII9664 | M | 60 | 23.13 | 227 | 61 | 166 |
| Mus*Dom | PWD | STRA | 23/06/2016 | ISII7891 | M | 60 | 18.62 | 251 | 33 | 218 |
| Mus*Dom | PWD | STRB | 04/08/2016 | ISII8449 | M | 60 | 20.77 | 146 | 15 | 131 |
| Mus*Dom | PWD | WLA | 19/01/2019 | EX2470 | M | 60 | 16.06 | 2 | 1 | 1 |
| Mus*Mus | PWD | BUSNA | 30/05/2018 | EX1130 | M | 60 | 16.74 | 235 | 31 | 204 |
| Mus*Mus | PWD | MPB | 19/06/2018 | EX1237 | M | 60 | 19.92 | 83 | 11 | 72 |
| Mus*Mus | PWD | PWD | 22/02/2016 | ISII6102 | M | 60 | 12.7 | 84 | 10 | 74 |
| Mus*Mus | PWD | SKA | 22/12/2017 | EX409 | M | 60 | 16.62 | 195 | 21 | 174 |
| Mus*Mus | PWD | STUF | 17/10/2016 | ISII9315 | M | 60 | 21.78 | 255 | 38 | 217 |
| Mus*Mus | PWD | STUS | 26/12/2016 | SY344 | M | 60 | 18.61 | 276 | 40 | 236 |
| Mus*Dom | SENK | WLA | 25/12/2017 | EX433 | M | 60 | 20.76 | 361 | 33 | 328 |
| Mus*Mus | SENK | PWD | 01/11/2017 | EX268 | M | 60 | 14.25 | 209 | 18 | 191 |
| Mus*DyM | SKA | BL6 | 23/08/2018 | EX1666 | M | 60 | 22.92 | 53 | 6 | 47 |
| Mus*DyM | SKA | SPLY | 30/11/2017 | EX342 | M | 60 | 19.23 | 136 | 20 | 116 |
| Mus*Dom | SKA | BIK | 21/09/2018 | EX1790 | M | 60 | 17.54 | 154 | 9 | 145 |
| Mus*Dom | SKA | DCP | 18/10/2018 | EX1950 | M | 60 | 18.41 | 178 | 15 | 163 |

| | | | | | | | | | | |
|---|---|---|---|---|---|---|---|---|---|---|
| Mus*Dom | SKA | SCHUNT | 23/05/2018 | EX1091 | M | 60 | 15.77 | 171 | 27 | 144 |
| Mus*Dom | SKA | SIT | 23/02/2018 | EX618 | M | 60 | 20.13 | 161 | 34 | 127 |
| Mus*Dom | SKA | STRA | 26/12/2017 | EX413 | M | 60 | 20.11 | 162 | 28 | 134 |
| Mus*Mus | SKA | BULS | 29/12/2017 | EX428 | M | 60 | 18.27 | 231 | 28 | 203 |
| Mus*Mus | SKA | MPB | 23/06/2018 | EX1287 | M | 60 | 17.18 | 253 | 31 | 222 |
| Mus*Mus | SKA | PWD | 23/01/2018 | EX510 | M | 60 | 16.77 | 156 | 28 | 128 |
| Mus*Mus | SKA | SENK | 29/04/2018 | EX919 | M | 60 | 15.61 | 299 | 44 | 255 |
| Mus*Mus | SKA | SKA | 23/05/2018 | EX1092 | M | 60 | 14.89 | 163 | 29 | 134 |
| Mus*Mus | SKA | STUF | 16/06/2018 | EX1229 | M | 60 | 18.58 | 308 | 31 | 277 |
| Mus*Mus | SKA | STUS | 16/03/2018 | EX701 | M | 60 | 13.92 | 169 | 18 | 151 |
| Mus*DyM | SKE | BL6 | 07/06/2018 | EX1179 | M | 60 | 24.25 | 7 | 2 | 5 |
| Mus*Dom | SKE | DGA | 14/11/2018 | EX2114 | M | 60 | 19.48 | 201 | 21 | 180 |
| Mus*Dom | SKE | DJO | 12/10/2018 | EX1925 | M | 60 | 22.4 | 0 | 0 | 0 |
| Mus*Dom | SKE | SIN | 29/03/2018 | EX743 | M | 60 | 18.77 | 0 | 0 | 0 |
| Mus*Dom | SKE | SIT | 07/09/2018 | EX1746 | M | 60 | 21.69 | 23 | 4 | 19 |
| Mus*Dom | SKE | WLA | 26/10/2018 | EX2007 | M | 60 | 18.91 | 30 | 5 | 25 |
| Mus*Mus | SKE | PWD | 03/01/2018 | EX432 | M | 60 | 16.05 | 155 | 15 | 140 |
| Mus*Mus | SKE | SKE | 25/05/2018 | EX1097 | M | 60 | 13.43 | 89 | 7 | 82 |
| Mus*Mus | SKE | STUS | 18/01/2018 | EX500 | M | 60 | 12.68 | 143 | 19 | 124 |
| Mus*DyM | STUF | SPLY | 22/07/2017 | SY3256 | M | 60 | 27.32 | 343 | 33 | 310 |
| Mus*Dom | STUF | DCP | 17/05/2018 | EX1057 | M | 60 | 25.87 | 299 | 23 | 276 |
| Mus*Dom | STUF | SCHUNT | 02/05/2017 | SY2006 | M | 60 | 24.13 | 362 | 52 | 310 |
| Mus*Dom | STUF | STRB | 18/06/2017 | SY2696 | M | 61 | 23.05 | 327 | 37 | 290 |
| Mus*Dom | STUF | WLA | 03/11/2018 | EX2070 | M | 60 | 22.12 | 284 | 26 | 258 |
| Mus*Mus | STUF | BULS | 07/06/2018 | EX1186 | M | 60 | 20.36 | 191 | 27 | 164 |
| Mus*Mus | STUF | STUF | 07/07/2016 | ISII8081 | M | 60 | 20.95 | 257 | 31 | 226 |
| Mus*Mus | STUF | STUS | 17/03/2016 | ISII6448 | M | 60 | 19.53 | 212 | 28 | 184 |
| Mus*DyM | STUS | SPLY | 06/01/2017 | SY456 | M | 60 | 20.67 | 160 | 31 | 129 |
| Mus*Dom | STUS | SIN | 28/05/2010 | SU6169 | M | 60 | 16.9 | 0 | 0 | 0 |
| Mus*Dom | STUS | STRB | 01/07/2017 | SY2956 | M | 60 | 18.99 | 78 | 20 | 58 |
| Mus*Mus | STUS | BULS | 17/06/2017 | SY2668 | M | 60 | 19.66 | 233 | 23 | 210 |
| Mus*Mus | STUS | PWD | 14/06/2017 | SY2601 | M | 60 | 17.13 | 187 | 19 | 168 |
| Mus*Mus | STUS | STUF | 20/12/2016 | SY227 | M | 60 | 17.86 | 269 | 64 | 205 |
| Mus*Mus | STUS | STUS | 30/01/2016 | ISII5891 | M | 60 | 12.63 | 160 | 30 | 130 |

| freq DSH | N Heads | N abnormal | N normal | freq abnormal |
|---|---|---|---|---|
| 6.37 | 193 | 18 | 175 | 9.326 |
| 5.82 | 227 | 28 | 199 | 12.335 |
| 13.1 | 175 | 8 | 167 | 4.571 |
| 10.03 | 200 | 9 | 191 | 4.5 |
| 10.61 | 188 | 20 | 168 | 10.638 |
| 14.87 | 176 | 25 | 151 | 14.205 |
| 33.33 | 6 | 3 | 3 | 50 |
| 7.69 | 77 | 10 | 67 | 12.987 |
| 5.06 | 141 | 11 | 130 | 7.801 |
| NA | 0 | 0 | 0 | NA |
| 6.95 | 223 | 10 | 213 | 4.484 |
| 18.91 | 126 | 7 | 119 | 5.556 |
| 10.32 | 157 | 10 | 147 | 6.369 |
| 14.22 | 136 | 9 | 127 | 6.618 |
| 5.19 | 188 | 9 | 179 | 4.787 |
| 7.81 | 149 | 9 | 140 | 6.04 |
| 12.45 | 159 | 12 | 147 | 7.547 |
| 9.04 | 119 | 8 | 111 | 6.723 |
| 12.5 | 122 | 15 | 107 | 12.295 |
| 10.05 | 110 | 8 | 102 | 7.273 |
| 4.76 | 55 | 8 | 47 | 14.545 |
| 11.85 | 195 | 15 | 180 | 7.692 |
| 9.27 | 126 | 6 | 120 | 4.762 |
| 15.73 | 40 | 4 | 36 | 10 |
| 13.1 | 135 | 5 | 130 | 3.704 |
| 3.7 | 83 | 13 | 70 | 15.663 |
| 6.35 | 34 | 8 | 26 | 23.529 |
| 9.73 | 110 | 9 | 101 | 8.182 |
| 6.62 | 76 | 8 | 68 | 10.526 |
| 11.63 | 51 | 7 | 44 | 13.725 |
| 3.64 | 141 | 12 | 129 | 8.511 |
| 10.74 | 74 | 8 | 66 | 10.811 |
| 3.7 | 105 | 15 | 90 | 14.286 |
| 7.31 | 130 | 8 | 122 | 6.154 |
| 6.1 | 135 | 14 | 121 | 10.37 |
| 6.02 | 108 | 8 | 100 | 7.407 |
| 15.95 | 100 | 8 | 92 | 8 |
| 5.29 | 127 | 9 | 118 | 7.087 |
| 6.77 | 109 | 15 | 94 | 13.761 |
| 6.3 | 143 | 12 | 131 | 8.392 |
| 6.76 | 127 | 13 | 114 | 10.236 |
| 6.76 | 117 | 10 | 107 | 8.547 |
| 8.18 | 130 | 12 | 118 | 9.231 |

| | | | | |
|---|---|---|---|---|
| 7.56 | 83 | 7 | 76 | 8.434 |
| 2.63 | 38 | 12 | 26 | 31.579 |
| 12.9 | 17 | 5 | 12 | 29.412 |
| 6.38 | 182 | 12 | 170 | 6.593 |
| 10.42 | 165 | 9 | 156 | 5.455 |
| 5.18 | 139 | 20 | 119 | 14.388 |
| 9.02 | 162 | 9 | 153 | 5.556 |
| 6.74 | 131 | 9 | 122 | 6.87 |
| 8.33 | 93 | 14 | 79 | 15.054 |
| 5.86 | 159 | 10 | 149 | 6.289 |
| 16.13 | 61 | 4 | 57 | 6.557 |
| 8.64 | 133 | 11 | 122 | 8.271 |
| 10.1 | 129 | 15 | 114 | 11.628 |
| 8.93 | 210 | 12 | 198 | 5.714 |
| 11.81 | 256 | 18 | 238 | 7.031 |
| 10.97 | 191 | 14 | 177 | 7.33 |
| 6.94 | 225 | 12 | 213 | 5.333 |
| 9.9 | 182 | 14 | 168 | 7.692 |
| 13.24 | 121 | 11 | 110 | 9.091 |
| 8.43 | 200 | 20 | 180 | 10 |
| 9.89 | 157 | 5 | 152 | 3.185 |
| 14.44 | 119 | 11 | 108 | 9.244 |
| 7.99 | 233 | 19 | 214 | 8.155 |
| 16.36 | 138 | 10 | 128 | 7.246 |
| 21.43 | 16 | 3 | 13 | 18.75 |
| 10.31 | 63 | 9 | 54 | 14.286 |
| 14.61 | 178 | 16 | 162 | 8.989 |
| 10.6 | 142 | 13 | 129 | 9.155 |
| 15.08 | 120 | 9 | 111 | 7.5 |
| 5.44 | 81 | 16 | 65 | 19.753 |
| 10.63 | 171 | 18 | 153 | 10.526 |
| 15.66 | 110 | 11 | 99 | 10 |
| 4.66 | 131 | 9 | 122 | 6.87 |
| 3.45 | 91 | 8 | 83 | 8.791 |
| 7.08 | 136 | 9 | 127 | 6.618 |
| 6.31 | 133 | 9 | 124 | 6.767 |
| 9.27 | 173 | 22 | 151 | 12.717 |
| 12.83 | 134 | 7 | 127 | 5.224 |
| 8.09 | 111 | 6 | 105 | 5.405 |
| 9.48 | 211 | 10 | 201 | 4.739 |
| 10.45 | 167 | 9 | 158 | 5.389 |
| 8.08 | 208 | 13 | 195 | 6.25 |
| 8.96 | 180 | 18 | 162 | 10 |
| 8.93 | 65 | 3 | 62 | 4.615 |
| 8.38 | 104 | 7 | 97 | 6.731 |
| 7.09 | 149 | 16 | 133 | 10.738 |
| 7.75 | 82 | 14 | 68 | 17.073 |
| 12.33 | 87 | 9 | 78 | 10.345 |
| 21.52 | 46 | 3 | 43 | 6.522 |
| 12.3 | 213 | 17 | 196 | 7.981 |

| | | | | |
|---|---|---|---|---|
| 13.45 | 102 | 5 | 97 | 4.899 |
| 12.77 | 81 | 12 | 69 | 14.815 |
| 10.71 | 36 | 17 | 19 | 47.222 |
| 8.06 | 44 | 14 | 30 | 31.818 |
| 10.61 | 114 | 6 | 108 | 5.263 |
| 11.8 | 238 | 16 | 222 | 6.723 |
| 8.59 | 186 | 12 | 174 | 6.452 |
| 9.7 | 199 | 10 | 189 | 5.025 |
| 6.52 | 211 | 20 | 191 | 9.479 |
| 10.98 | 207 | 17 | 190 | 8.213 |
| 11.34 | 209 | 11 | 198 | 5.263 |
| 13.57 | 130 | 13 | 117 | 10 |
| 6.86 | 97 | 14 | 83 | 14.433 |
| 10.12 | 98 | 7 | 91 | 7.143 |
| 8.91 | 191 | 7 | 184 | 3.665 |
| 7.24 | 122 | 14 | 108 | 11.475 |
| 8.23 | 90 | 13 | 77 | 14.444 |
| 13.1 | 95 | 7 | 88 | 7.368 |
| 10.71 | 146 | 9 | 137 | 6.164 |
| 14.49 | 133 | 23 | 110 | 17.293 |
| 11.02 | 226 | 18 | 208 | 7.965 |
| 7.81 | 194 | 13 | 181 | 6.701 |
| 6.56 | 208 | 13 | 195 | 6.25 |
| 17.73 | 172 | 13 | 159 | 7.558 |
| 13.21 | 198 | 17 | 181 | 8.586 |
| 12.76 | 122 | 10 | 112 | 8.197 |
| 5.92 | 107 | 9 | 98 | 8.411 |
| 5.65 | 150 | 19 | 131 | 12.667 |
| 5.3 | 145 | 13 | 132 | 8.966 |
| 9.34 | 140 | 8 | 132 | 5.714 |
| 7.82 | 169 | 10 | 159 | 5.917 |
| 12.37 | 64 | 10 | 54 | 15.625 |
| 9.96 | 166 | 7 | 159 | 4.217 |
| 5.88 | 109 | 11 | 98 | 10.092 |
| 8.33 | 134 | 11 | 123 | 8.209 |
| 9.89 | 158 | 13 | 145 | 8.228 |
| 12.82 | 75 | 11 | 64 | 14.667 |
| 16.55 | 83 | 4 | 79 | 4.819 |
| 14.05 | 110 | 9 | 101 | 8.182 |
| 17.11 | 155 | 11 | 144 | 7.097 |
| 25 | 12 | 7 | 5 | 58.333 |
| 42.86 | 7 | 3 | 4 | 42.857 |
| 20.29 | 136 | 11 | 125 | 8.088 |
| 22.28 | 123 | 13 | 110 | 10.569 |
| 12.04 | 71 | 15 | 56 | 21.127 |
| 8.18 | 97 | 10 | 87 | 10.309 |
| 5.29 | 106 | 5 | 101 | 4.717 |
| 12.2 | 92 | 6 | 86 | 6.522 |
| 17.56 | 129 | 10 | 119 | 7.752 |
| 5.15 | 85 | 5 | 80 | 5.882 |

| | | | | |
|---|---|---|---|---|
| 19.01 | 82 | 2 | 80 | 2.439 |
| 8.73 | 224 | 11 | 213 | 4.911 |
| 23.08 | 61 | 9 | 52 | 14.754 |
| 13.57 | 200 | 14 | 186 | 7 |
| 18.25 | 163 | 13 | 150 | 7.975 |
| 15.23 | 167 | 13 | 154 | 7.784 |
| 13.49 | 160 | 13 | 147 | 8.125 |
| 8.09 | 179 | 12 | 167 | 6.704 |
| 12.81 | 150 | 10 | 140 | 6.667 |
| 9.5 | 130 | 15 | 115 | 11.538 |
| 14.05 | 116 | 8 | 108 | 6.897 |
| 8.3 | 153 | 16 | 137 | 10.458 |
| 9.8 | 139 | 9 | 130 | 6.475 |
| 15.04 | 79 | 5 | 74 | 6.329 |
| 15.04 | 169 | 10 | 159 | 5.917 |
| 9.85 | 143 | 6 | 137 | 4.196 |
| 9.58 | 141 | 11 | 130 | 7.801 |
| 8.75 | 194 | 9 | 185 | 4.639 |
| 13.39 | 82 | 5 | 77 | 6.098 |
| 9.96 | 147 | 14 | 133 | 9.524 |
| 14.04 | 164 | 11 | 153 | 6.707 |
| 15.7 | 105 | 9 | 96 | 8.571 |
| 15.48 | 96 | 12 | 84 | 12.5 |
| 12.6 | 68 | 6 | 62 | 8.824 |
| 10.92 | 176 | 8 | 168 | 4.545 |
| 4.84 | 74 | 4 | 70 | 5.405 |
| 9.18 | 119 | 10 | 109 | 8.403 |
| NA | 0 | 0 | 0 | NA |
| 17.58 | 102 | 7 | 95 | 6.863 |
| 11.64 | 141 | 13 | 128 | 9.22 |
| 12.39 | 135 | 12 | 123 | 8.889 |
| 28.57 | 14 | 9 | 5 | 64.286 |
| 15.45 | 150 | 14 | 136 | 9.333 |
| 20.51 | 162 | 12 | 150 | 7.407 |
| 26.87 | 135 | 7 | 128 | 5.185 |
| 13.15 | 147 | 12 | 135 | 8.163 |
| 10.27 | 83 | 8 | 75 | 9.639 |
| 50 | 0 | 0 | 0 | NA |
| 13.19 | 136 | 6 | 130 | 4.412 |
| 13.25 | 50 | 1 | 49 | 2 |
| 11.9 | 55 | 4 | 51 | 7.273 |
| 10.77 | 118 | 10 | 108 | 8.475 |
| 14.9 | 159 | 16 | 143 | 10.063 |
| 14.49 | 173 | 21 | 152 | 12.139 |
| 9.14 | 218 | 12 | 206 | 5.505 |
| 8.61 | 129 | 9 | 120 | 6.977 |
| 11.32 | 31 | 4 | 27 | 12.903 |
| 14.71 | 81 | 10 | 71 | 12.346 |
| 5.84 | 88 | 9 | 79 | 10.227 |
| 8.43 | 96 | 5 | 91 | 5.208 |

| | | | | |
|---|---|---|---|---|
| 15.79 | 105 | 8 | 97 | 7.619 |
| 21.12 | 100 | 11 | 89 | 11 |
| 17.28 | 89 | 10 | 79 | 11.236 |
| 12.12 | 138 | 9 | 129 | 6.522 |
| 12.25 | 154 | 9 | 145 | 5.844 |
| 17.95 | 100 | 9 | 91 | 9 |
| 14.72 | 173 | 8 | 165 | 4.624 |
| 17.79 | 96 | 7 | 89 | 7.292 |
| 10.06 | 173 | 6 | 167 | 3.468 |
| 10.65 | 115 | 8 | 107 | 6.957 |
| 28.57 | 7 | 5 | 2 | 71.429 |
| 10.45 | 122 | 9 | 113 | 7.377 |
| NA | 0 | 0 | 0 | NA |
| NA | 0 | 0 | 0 | NA |
| 17.39 | 23 | 18 | 5 | 78.261 |
| 16.67 | 20 | 12 | 8 | 60 |
| 9.68 | 91 | 7 | 84 | 7.692 |
| 7.87 | 64 | 7 | 57 | 10.938 |
| 13.29 | 94 | 9 | 85 | 9.574 |
| 9.62 | 218 | 12 | 206 | 5.505 |
| 7.69 | 185 | 10 | 175 | 5.405 |
| 14.36 | 210 | 11 | 199 | 5.238 |
| 11.31 | 175 | 10 | 165 | 5.714 |
| 9.15 | 164 | 11 | 153 | 6.707 |
| 14.14 | 121 | 10 | 111 | 8.264 |
| 12.06 | 151 | 9 | 142 | 5.96 |
| 13.21 | 118 | 16 | 102 | 13.559 |
| 19.38 | 98 | 22 | 76 | 22.449 |
| NA | 0 | 0 | 0 | NA |
| 25.64 | 52 | 10 | 42 | 19.231 |
| 9.87 | 148 | 9 | 139 | 6.081 |
| 10.16 | 109 | 6 | 103 | 5.505 |
| 23.79 | 164 | 23 | 141 | 14.024 |
| 18.75 | 96 | 14 | 82 | 14.583 |
