## Supplementary Table 2.1-2.4 for "Sperm quality parameters are increased and asymmetric in house mouse hybrids"

**Supplementary Table S2.1** Medians and the 25 and 75% quartiles for abnormal head sperm (ASH) and dissociated sperm heads (DSH) in intra- and intersubspecific crosses

| Cross_type | Medians.ASH | Medians.DSH |
|---|---|---|
| Dom*Dom | 7.10 ( 5.35 ~ 8.44 ) | 9.93 ( 8.11 ~ 13.03 ) |
| Dom*Mus | 10.00 ( 7.14 ~ 14.39 ) | 8.33 ( 6.62 ~ 10.71 ) |
| Mus*Dom | 8.13 ( 6.53 ~ 10.48 ) | 15.13 ( 10.66 ~ 18.04 ) |
| Mus*Mus | 6.98 ( 5.84 ~ 9.00 ) | 12.06 ( 9.68 ~ 14.05 ) |

**Supplementary Table S2.2**

**Dissociated sperm heads tests**

**(a) anova for linear model based on Box-Cox transformed data**

formula: DSH.BC ~ OffCross

| | Df | Sum Sq | Mean Sq | F value | Pr(>F) |
|---|---|---|---|---|---|
| OffCross | 3 | 5.483 | 1.828 | 18.413 | 2.019e-10 |
| Residuals | 174 | 17.271 | 0.099 | | |

**(b) anova for mixed effects model based on Box-Cox transformed data**

Formula: DSH.BC ~ OffCross + (1 | StrSir) + (1 | StrDam)

| | Sum Sq | Mean Sq | NumDF | DenDF | F value | Pr(>F) |
|---|---|---|---|---|---|---|
| OffCross | 2.189 | 0.730 | 3 | 44.927 | 9.682 | 4.799e-05 |

**(c) comparison linear and mixed effects model**

Models:

lmDSH: DSH.BC ~ OffCross

mixDSH.MF: DSH.BC ~ OffCross + (1 | StrSir) + (1 | StrDam)

| | Df | AIC | BIC | logLik | deviance | Chisq | Chi Df | Pr(>Chisq) |
|---|---|---|---|---|---|---|---|---|
| lmDSH | 5 | 99.914 | 115.82 | -44.957 | 89.914 | | | |
| mixDSH.MF | 7 | 93.623 | 115.90 | -39.812 | 79.623 | 10.291 | 2 | 0.005826 |

ΔAIC = 6.29

**Supplementary Table S2.3** Results of *post hoc* pairwise comparisons of dissociated sperm heads (DSH) between cross types using Tukey contrasts. Analysis is based on the linear mixed model with random effects using transformed data

| Contrast | Estimate±SE | Prob |
|---|---|---|
| Dom*Mus - Dom*Dom | -0.105 ± 0.081 | 0.550 |
| Mus*Dom - Dom*Dom | 0.354 ± 0.082 | <0.001 |
| Mus*Mus - Dom*Dom | 0.144 ± 0.099 | 0.452 |
| Mus*Dom - Dom*Mus | 0.459 ± 0.091 | <0.001 |
| Mus*Mus - Dom*Mus | 0.249 ± 0.074 | 0.004 |
| Mus*Mus - Mus*Dom | 0.210 ± 0.083 | 0.052 |

**Supplementary Table S2.4** Results of a nonparametric Tukey-Kramer (Nemenyi) test for *post hoc* pairwise comparisons of abnormal sperm heads (ASH) between cross types

| Contrast | Estimate | Prob |
|---|---|---|
| Dom*Mus - Dom*Dom | 5.903 | 0.000 |
| Mus*Dom - Dom*Dom | 3.347 | 0.084 |
| Mus*Mus - Dom*Dom | 0.988 | 0.898 |
| Mus*Dom - Dom*Mus | 2.450 | 0.307 |
| Mus*Mus - Dom*Mus | 5.121 | 0.002 |
| Mus*Mus - Mus*Dom | 2.473 | 0.298 |
